# Phylogeny of the *Formicoxenus* genus-group (Hymenoptera: Formicidae) reveals isolated lineages of *Leptothorax acervorum* in the Iberian Peninsula predating the Last Glacial Maximum

**DOI:** 10.1101/2021.11.05.467305

**Authors:** Dario I. Ojeda, Max John, Robert L. Hammond, Riitta Savolainen, Kari Vepsäläinen, Torstein Kvamme

## Abstract

The *Formicoxenus* genus-group comprises six genera within the tribe Crematogastrini. The group is well known for repeated evolution of social parasitism among closely related taxa and cold-adapted species with large distribution ranges in the Nearctic and Palearctic regions. Previous analyses based on nuclear markers (ultraconserved elements, UCEs) and mitochondrial genes suggest close relationship between *Formicoxenus* Mayr, 1855, *Leptothorax* Mayr, 1855 and *Harpagoxenus* Forel, 1893. However, scant sampling has limited phylogenetic assessment of these genera. Also, previous phylogeographic analyses of *L. acervorum* (Fabricius, 1793) have been limited to its West-Palearctic range of distribution, which has provided a narrow view on recolonization, population structure and existing refugia of the species. Here, we inferred the phylogenenetic history of genera within the *Formicoxenus* genus-group and reconstructed the phylogeography of *L. acervorum* with more extensive sampling. We employed four datasets consisting of whole genomes and sequences of the COI. The topologies of previous nuclear and our inferences based on mitochondrial genomes were overall congruent. Further, *Formicoxenus* may not be monophyletic. We found several monophyletic lineages that do not correspond to the current species described within *Leptothorax*, especially in the Nearctic region. We identified a monophyletic *L. acervorum* lineage that comprises both Nearctic and Palearctic locations. The most recent expansion within *L. acervorum* probably occurred within the last 0.5 Ma with isolated populations predating the Last Glacial Maximum (LGM), which are localized in at least two refugial areas (Pyrenean and Northern plateau) in the Iberian Peninsula. The patterns recovered suggest a shared glacial refugium in the Iberian Peninsula with cold-adapted trees that currently share high-altitude environments in this region.

## 1. Introduction

Several invertebrate groups have species with Holarctic distribution, including beetles (Larson and Nilsson, 1985), Lepidoptera (Landry et al., 2013), spiders (Marusik and Koponen, 2005) and ants (Schär et al., 2018). Among ant species, only three species (*Camponotus herculeanus* Linnaeus, 1758, *Formica gagatoides* Ruzsky, 1904 and *Leptothorax acervorum* Fabricius, 1793) are known to have monophyletic lineages with a Holarctic distribution (Schär et al., 2018). The latter of these ant species belongs to the tribe Crematogastrini Emery, 1914 within the subfamily Myrmicinae, with Crematogastrini comprising some 6,630 species (Blaimer et al., 2018; Ward et al., 2015). There, recent phylogenomic analyses based on ultraconserved elements (UCEs) have consistently recovered a monophyletic lineage of six genera (*Vombisidris* Bolton 1991, *Gauromyrmex* Menozzi, 1993, *Harpagoxenus* Forel, 1893, *Formicoxenus* Mayr, 1855, *Temnothorax* Mayr, 1861 and *Leptothorax* Mayr, 1855) referred to informally as the *Formicoxenus* genus-group (Blaimer et al., 2018; Branstetter et al., 2017). These analyses have recovered a close relationship between *Formicoxenus* and *Leptothorax*. However, all these studies have been focused on higher taxonomic relationships and few studies have included a comprehensive sampling of species within each genus (Heinze and Gratiashvili, 2015; Prebus, 2017; Schär et al., 2018). Among the *Formicoxenus* genus-group, *Leptothorax* is the second largest genus with an estimated 20 species (AntWeb ver. 8.42, https://www.antweb.org, accessed 29 October 2020). The genus has a Holarctic distribution and it has been inferred to have originated in the Nearctic and dispersed in the Palearctic within the last 2 Ma (Schär et al., 2018). Relationships within *Leptothorax* have not been fully resolved and previous phylogenetic analyses indicate the presence of multiple undescribed and non-monophyletic taxa, particularly in the Nearctic (Heinze and Gratiashvili, 2015; Schär et al., 2018). At least seven species have been described in the Palearctic region, most of them with limited distribution and only *L. acervorum* with a distribution both in the Nearctic and the Palearctic regions (Schär et al., 2018).

Taxa that span large geographic regions in both the Nearctic and Palearctic are expected to have experienced variable connectivity because of the fluctuating presence of the land bridge of Beringia between Eurasia and North America (DeChaine, 2008). Also, climate oscillations during the Quaternary (last 2 Ma), characterized by pronounced cycles between cold glacial (ca. 100,000 years) and warm interglacial periods (ca. 20,000 years) during the last 700,000 years, altered the geographic distribution of species in the northern hemisphere (Nearctic and Palearctic) (Hewitt, 2000). During these glacial episodes, species ranges contracted to refugia in suitable areas in the southern part of their distribution. As the climate warmed and glaciers retreated, species with a temperate range of distribution expanded and reconnected. In contrast, the range of distribution for boreal cold-adapted species likely reduced and fragmented (Hewitt, 1996; Petit et al., 2003). Evidence from several ant species have suggested the presence of several refugia during the Pleistocene (2.58 - 0.012 Ma) in the southern Mediterranean peninsulas, the Caspian-Caucasus region and further east in southern East Asia (Beibl et al., 2007; Goropashnaya et al., 2004; Leppänen et al., 2013, 2011; Pusch et al., 2006; Schlick-Steiner et al., 2007). In addition, populations of cold-adapted ant species could have also survived in more northerly refugia near the permafrost (Leppänen et al., 2013, 2011). Indeed, *L. acervorum* is among the very few cold-adapted species that extend their distribution above the polar circle both in the Nearctic and the Palearctic (Berman et al., 2010; Heinze et al., 1998, 1996). In the Palearctic, this species occurs in the boreal zone from the Atlantic Ocean to Japan, and in the mountains of southern Europe, the Caucasus, and the Tien-Shan and Pamir (Czechowski et al., 2012; Seifert, 2018). Thus, the climatic fluctuations of the Quaternary have likely played a significant role in shaping its current distribution, connectivity, and genetic diversity. Populations located near the permafrost and those located on the southern range of its distribution were likely affected differently.

The most recent phylogenetic analysis of *Leptothorax* indicate that *L. acervorum* originated about 2 Ma, with the least diverged populations located in the Nearctic region. Within the Palearctic, populations situated in the Iberian Peninsula were inferred to be the less divergent among the specimens included (Schär et al., 2018), which might have been located in refugia during the glacial cycles. In addition, more detailed analyses based on mitochondrial DNA (COI-3P region) and microsatellites (SSRs) have been used to infer the phylogeography and population structure of this species in the western part of its distribution (West Palearctic). These analyses have found generally less population structure in *L. acervorum* compared to other species closely related species within *Leptothorax*, e.g. *L. muscorum* (Nylander, 1846) and *Harpagoxenus sublaevis* (Nylander, 1849) (Brandt et al., 2007; Foitzik et al., 2009; Trettin et al., 2016), but also evidence of divergent haplotypes have been found in the Pyrenees and Southern France (Trettin et al., 2016). Given the large distribution range across the Holarctic and the extensive variation in the latitudinal range in western Europe (from the Iberian Peninsula to North Cape in Norway) (Heinze and Holldobler, 1994), approaches that combine analyses at different taxonomic levels with extensive sampling are necessary to understand the phylogenetic relationships and evolutionary history of *Leptothorax* species. Here we present the most comprehensive sampling of members of the *Formicoxenus* genus-group with an emphasis on the phylogenetic relationships within *Leptothorax* and the biogeography of *L. acervorum* across its range of distribution in the Holarctic region. Our specific objectives are: 1) to infer relationships among the six genera of the *Formicoxenus* genus-group using whole mitochondrial genomes and asses their correspondence with previous topologies obtained with nuclear genes, 2) to clarify the relationships of *Leptothorax* species and the timing of divergence of the Palearctic species, 3) to determine if the populations of *L. acervorum* situated in the Iberian Peninsula survived in different refugia during the glacial cycles of the Quaternary.

## 2. Materials and methods

### 2.1 Taxon sampling and datasets

The sampling strategy used in this study was developed to represent the *Formicoxenus* genus-group at three different hierarchical levels. The first dataset consisted of 49 specimens representing all six genera (*Temnothorax, Leptothorax, Formicoxenus, Harpagoxenus, Gauromyrmex* and *Vombisidris*) currently recognized within this group (Blaimer et al., 2015; Prebus, 2017), representing 17% of genera within the Crematogastrini. We also included outgroups from Myrmicinae (all tribes), Dolichoderinae and Ponerinae. In this data set we used whole mitochondrial genomes to explore the major relationships within *Formicoxenus* genus-group at the genus level. We used only one representative specimen per species within each genus, except for *L. acervorum*, where we included multiple samples (Table S1). In the second dataset, we gathered specimens representing eight out of the 20 *Leptothorax* species currently recognized (AntWeb ver. 8.42, https://www.antweb.org, accessed 29 October 2020), two *Formicoxenus* species and *H. sublaevis*. In this dataset we sequenced the section of the mitochondrial cytochrome c oxidase (COI-5P region, 658 bp) in 96 specimens (Table S2). The third dataset consisted of 113 specimens of *L. acervorum* across its distribution range in the Holarctic region, where we sequenced the same gene region (COI-5P region, 658 bp) as the previous dataset. This dataset was complemented with available sequences from public repositories (Table S3). Finally, dataset four consisted of the mitochondrial region COI-*trnL2*-COII (2,293 bp) and it was obtained from a total of 80 *L. acervorum* specimens from six populations in Spain and the UK (Table S3).

### 2.2. Whole mitochondrial sequencing and assembly

Mitochondrial genomes were newly generated for six specimens of *L. acervorum* from six different populations in Spain and the UK (Spain: Valdelinares (V), Orihuela del Tremedal (OT), Larra (L), Niela Refuge (NR), Pla de la Font (PF); UK: Santon Downham (SD) (Table S3). A *de novo* mitochondrial genome was identified as part of a whole genome sequencing project from a single adult male (PF population, sample: PF18_15_M1) using 10x linked reads assembled with Supernova 2.1.1 (Weisenfeld et al., 2018). The scaffold containing the mtDNA genome was identified by a BLASTn query of the assembled genome with two published *L. acervorum* mtDNA sequences (query 1: COXI – tRNA - Leu - COXII: GenBank: KU245569 (Trettin et al., 2016); query 2: COB: GenBank: HQ259995 (Gill et al., 2009). These two sequences, located ∼6Kb apart in the canonical hymenopteran mtDNA genome, were used to minimize erroneous matches to nuclear genomic scaffolds containing translocated mtDNA (NUMTs). Only two scaffolds (102,807 and 104,071) showed convincing matches to both query sequences (E value = 0, bit scores > 1000). However, mapping re-sequenced samples (see below) showed scaffold 102,807 had 40 times higher coverage (200x-400x) than 104,071 (∼5x-10x coverage) with the latter having similar coverage to the rest of the presumed nuclear genome. Furthermore, scaffold 102,807 was 17Kb in length (the expected size of the mtDNA genome) whereas scaffold 104,071 was longer than expected at 24Kb. These lines of evidence clearly show scaffold 102,807 contains the *L. acervorum* mtDNA genome whereas scaffold 104,071 is a transposition of mtDNA sequences to the nuclear genome (a NUMT).

To genotype single individuals in the six populations (V, OT, L, NR, PF, and SD), short-read sequence data (Illumina HiSeq 2×150bp paired-end reads) were, after quality control steps, aligned to the draft genome with Bowtie2 2.3.5 (Langmead and Salzberg, 2014) and processed with SAMtools (Li et al., 2009) to produce bam files. Bam files were then subset to only include the identified mtDNA scaffold (scaffold: 102,807) with SAMtools. These mtDNA alignments were converted to mpileup with BCFtools (--max-depth 1000) and BCFtools call used to produce vcf files. Vcf files were indexed and normalized and variants within 5bp of any indels removed with BCFtools. Finally, a fasta file for each alignment was produced with BCFtools consensus (Supplementary Information, Online Methods).

In addition, mitochondrial genomes of the taxa within the *Formicoxenus* genus-group were extracted and assembled from ultra-conserved elements (UCE) libraries from previous studies (Branstetter et al., 2017; Prebus, 2017) using MitoFinder (Allio et al., 2020). Outgroup species within subfamilies Myrmicinae, Dolichoderinae and Ponerinae were downloaded from Genbank, previously published in several studies (Cicconardi et al., 2020; Du et al., 2019; Duan et al., 2016; Gotzek et al., 2010; Hasegawa et al., 2011; Liu et al., 2016; Park et al., 2021, 2020b, 2020a, 2019; Rodovalho et al., 2014). SRA sequences and the assembled mitochondrial genomes of *L. acervorum* are deposited in the NCBI Bioproject PRJNA634471.

### 2.3 Phylogenomic analyses using whole mitochondrial genomes

All 49 whole mitochondrial genomes were aligned using MAFFT ver. 7.310 (Katoh and Kuma, 2002) with default parameters. Visual inspection and further adjustment were performed with AliView (Larsson, 2014) and summary statistics of the alignment were obtained with AMAS (Borowiec, 2016). Phylogenetic analysis was performed with maximum likelihood (ML) as implemented in IQ-TREE 1.6.1 (Nguyen et al., 2015) with ultrafast likelihood bootstrap with 1000 replicates. The final tree was visualized and edited with FigTree (Rambaut, 2016).

### 2.4 DNA extraction, PCR amplification and sequencing of cytochrome c oxidase (COI)

We collected either pupae or adults of workers, males, or queens from different colonies of *H. sublaevis, L. acervorum, L. kutteri* (Buschinger, 1965) and *L. muscorum* (Table S2). DNA was extracted from legs or whole specimens using the salt extraction method (Aljanabi and Martinez, 1997) and we sequenced the portion of the mitochondrial COI using previous primers and PCR conditions (Folmer et al., 1994). The sequences obtained were edited, visually inspected using Sequencher (Gene Codes), and aligned with AliView (Larsson, 2014). COI sequences are deposited in the NCBI Bioproject PRJNA634471, under accessions XXXXXXXX-XXXXXXXX.

### 2.5 *Phylogenetic analyses and dating estimation within* Formicoxenus-Leptothorax *based on the COI gene region*

To infer the phylogenetic relationships within the *Formicoxenus-Leptothorax*, we used the 5’ region of COI (658 bp, ranging from 5442-6601 in the *L. acervorum* mitochondrial genome assembly). First, we performed an explorative analysis based on a comprehensive sampling from this gene region using the sequences generated in this study and available sequences from GenBank and BOLD: The Barcode of Life Data System (www.barcodinglife.org). The matrix was aligned and manually edited with AliView (Larsson, 2014), with summary statistics obtained with AMAS (Borowiec, 2016). This preliminary analysis was based in a total of 747 specimens of *Formicoxenus* (2 spp.), *Leptothorax* (8 spp.) and specimens of *H. sublaevis* (outgroup). We used maximum likelihood (ML) as implemented in IQ-TREE 1.6.1 (Nguyen et al., 2015) with ultrafast likelihood bootstrap with 1000 replicates (Minh et al., 2013). Based on the results from this analysis (data not shown), we selected representative specimens of the major Nearctic lineages identified (>80 bootstrap support) and all the specimens in the Palearctic lineage of *L. acervorum*. This final dataset consisted of 96 specimens representing eight *Leptothorax* spp., specimens labelled as *Leptothorax* sp., *L. muscorum* complex, *Leptothorax* sp. AF CAN, two *Formicoxenus* spp. and *H. sublaevis* as an outgroup. This dataset consisted of the newly generated sequences in this study and available sequences from previous publications (Hebert et al., 2016; Prebus, 2017; Schär et al., 2018; Smith et al., 2009; Stahlhut et al., 2013) (Table S2). The best nucleotide substitution model and ML analysis were inferred with IQ-TREE 1.6.1. Clade support was assessed with ultrafast likelihood bootstrap with 1000 replicates. In addition, we also performed a Bayesian inference (BI) as implemented in MrBayes 3.2.6 (Huelsenbeck and Ronquist, 2001; Ronquist et al., 2012) with four chains, two runs of 20 million generations with the invgamma rate of variation, the GTR+Г model of nucleotide substitution and a sample frequency of 1000. We used Tracer 1.7 (Rambaut et al., 2018) to verify whether effective samples sizes (ESS values) were higher than 200 for all parameters.

To estimate divergence times among the lineages in *Formicoxenus-Leptothorax*, we used a simplified dataset representing the same number of species as above, but fewer specimens (63) of these two genera. We used Beast 1.10.4 (Bouckaert et al., 2014; Suchard et al., 2018) with a strict clock model and a constant population size under a coalescence model. We employed the divergence time estimated in the Formicinae (Blaimer et al., 2015) by placing a prior in the divergence estimate of *Harpagoxenus* and *Formicoxenus*-*Leptothorax* of 8.89 (13.89-3.89) Ma. We ran two independent runs of 50 million generations each, sampling values every 1,000 steps. Output files were analyzed with Tracer 1.7 to assess chain convergence and LogCombiner 1.10.4 was used to combine independent runs. Finally, we used Treeannotator 1.10.4 to generate the maximum-clade-credibility tree. ML and BI consensus trees were visualized and edited with FigTree (Rambaut, 2016).

### 2.6 *Phylogeography and genetic diversity of* Leptothorax acervorum

To gain further insights into the geographic distribution of genetic diversity of *L. acervorum* across its Holarctic distribution, we first determined the number of haplotypes, haplotype diversity (*Hd*) and defined haplotypes with DnaSP ver. 6.12 (Rozas et al., 2017). Then, we reconstructed the haplotype network of all 113 specimens (Table S3) using the COI-5P gene region (647 bp, dataset 3) with the statistical parsimony network using TCS (Clement et al., 2002) as implemented in popart ver. 1.7 (Leigh and Bryant, 2015). Given that our interest was focused on the populations distributed in the Iberian Peninsula, we explored in more detail five populations from this region and one population from England (Table S3). In these analyses we used the mitochondrial region encompassing COI, *trnL2* and COII comprising 2,293 bp (dataset 4). Haplotype (*h*), polymorphic sites and nucleotide diversities (π) were calculated using the program DnaSP ver. 6.12. We employed Fu’s *Fs* and Tajima’s D tests of selective neutrality to determine whether *L. acervorum* populations from the Iberian Peninsula and England could have experienced recent expansions. Fu’s *F* is based in the infinite-site model and a population expansion increase of rare alleles in the population, leading to negative values (Fu, 1997). Tajima’s D uses the frequency of segregating nucleotide sites and the average number of nucleotide differences obtained from pairwise comparisons (Tajima, 1989). Deviations from neutrality could indicate the effect of selection and/or population size changes. Population expansions will increase rare alleles in the population, leading to values < 0, while population contractions (bottlenecks) will increase intermediate variants in the population. COI, *trnL2* and COII sequences are deposited in the NCBI Bioproject PRJNA634471, under accessions XXXXXXXX-XXXXXXXX.

## 3. Results

### 3.1. *Phylogenomic analyses of the* Formicoxenus *genus-group*

The final alignment of the mitochondrial genomes consisted of 14,351 bp with 13.32 % missing data, 69% of sites variable and 59% of sites parsimony informative (Supplementary Information, Data S1). We recovered monophyletic lineages for all the tribes, except Attini Smith, 1858 within Myrmicinae, with most branches having moderate (>75%) to high (>85%) bootstrap support. Our phylogenetic analysis recovered all six genera of the *Formicoxenus* genus-group as a monophyletic lineages within Crematogastrini, with *Formicoxenus* as the most closely related genus to *Leptothorax*. The most closely related tribe was Solenopsidini Forel, 1893 (Fig.1).

**Fig. 1.**
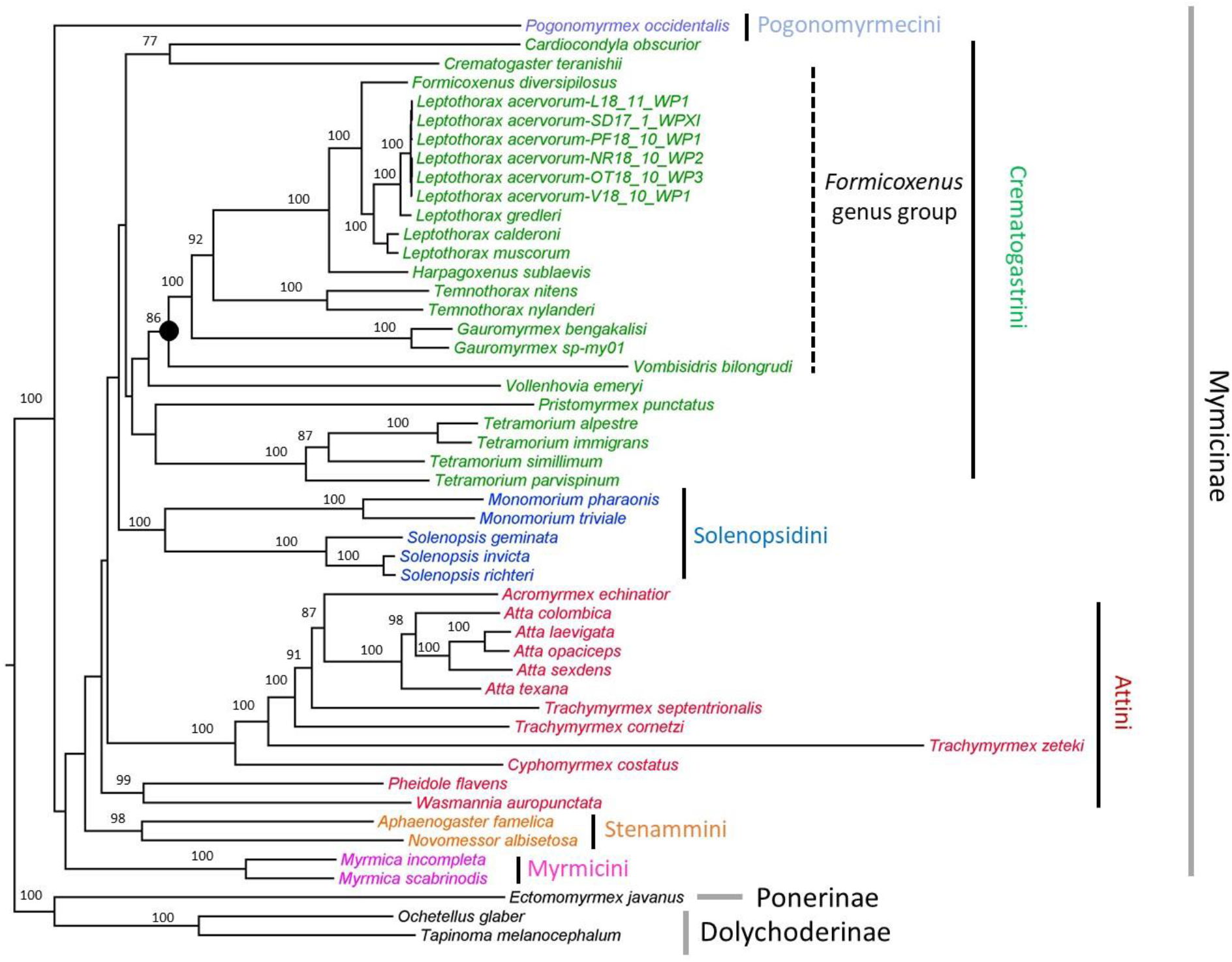
Best tree obtained in the phylogenetic analysis inferred with whole mitochondrial genomes of the *Formicoxenus* genus-group using ML as implemented in IQ-TREE. Values next to the branches represent bootstrap support (1000 bs replicates). Only branches with support over 70 are labelled. The black circle indicates the node with the genera of the *Formicoxenus* genus-group. Different colors indicate tribes within the Myrmycinae.

### 3.2 *Relationships within* Leptothorax *and divergence times of the Palearctic species*

The alignment matrix of the COI-5P region (dataset 2) consisted of 658 bp with 200 variable sites (30.4 %), 148 parsimony informative sites (22.5%) and 1.42% of missing data. We recovered the species of *Formicoxenus* on different lineages within *Leptothorax*, suggesting that the former genus might not represent a monophyletic lineage. All the three Palearctic *Leptothorax* species we included (*L. muscorum, L. gredleri* Mayr, 1855 and *L. kutteri*) represent monophyletic lineages, whereas specimens assigned to *L. muscorum* from the Nearctic region represent several undescribed taxa (Fig. 2 and Fig. S1). Similarly, we found non-monophyletic lineages for the other Nearctic species *L. canadensis* Provancher, 1887 and *L. calderoni* Creighton, 1950, but not for *L. retractus* Francoeur, 1986. Our divergence estimate suggests that the stem age lineages of Palearctic taxa (*L. gredleri, L. muscorum, L. kutteri* and *L. acervorum*) ranges between 1-1.6 Ma (Fig. 3). The crown age of the Palearctic lineage of *L. acervorum* was estimated at 0.56 Ma, with specimens from the Iberian Peninsula ranging in age between 0.1 and 0.5 Ma. The most recent derived lineage (0.30 Ma) within *L. acervorum* comprises both specimens from the Nearctic and Palearctic distribution, including specimens at high latitudes mainly from the West Nearctic (Fig. 4).

**Fig. 2.**
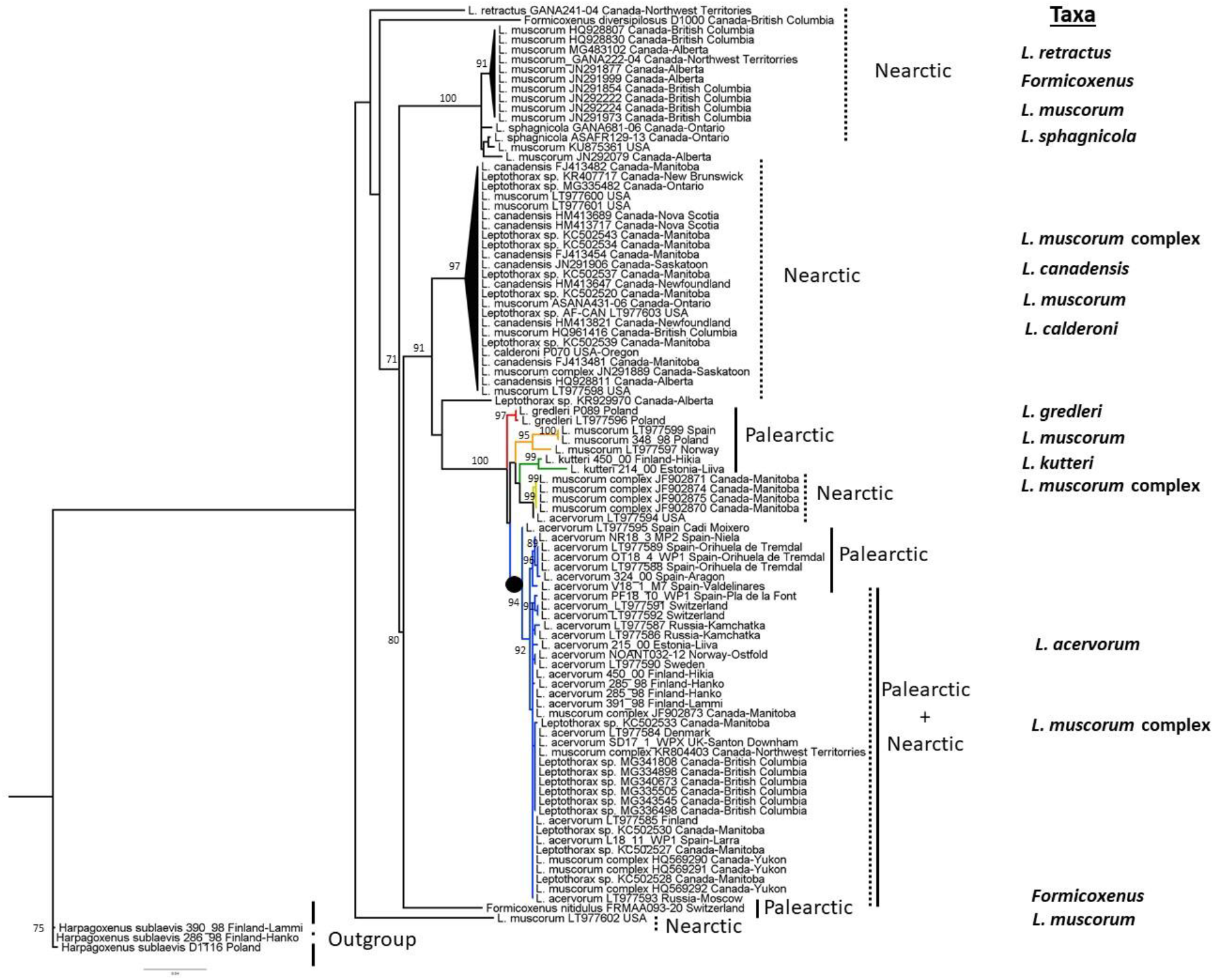
Best tree recovered of the analysis of *Formicoxenus*-*Leptothorax* using ML as implemented in IQ-TREE using the dataset of COI-5P region. The different colours of the branches indicate monophyletic lineages recovered on the species with Palearctic distribution. The black circle indicates the monophyletic lineage of *L. acervorum* (plus specimens of *L. muscorum* complex and *Leptothorax* sp.) that were later used in the phylogeographic analysis. Species currently recognized in these genera are indicated next to the lineages (taxa). Values next to the branches represent bootstrap support (1000 bs replicates). Only branches with >70 bootstrap support are labelled.

**Fig. 3.**
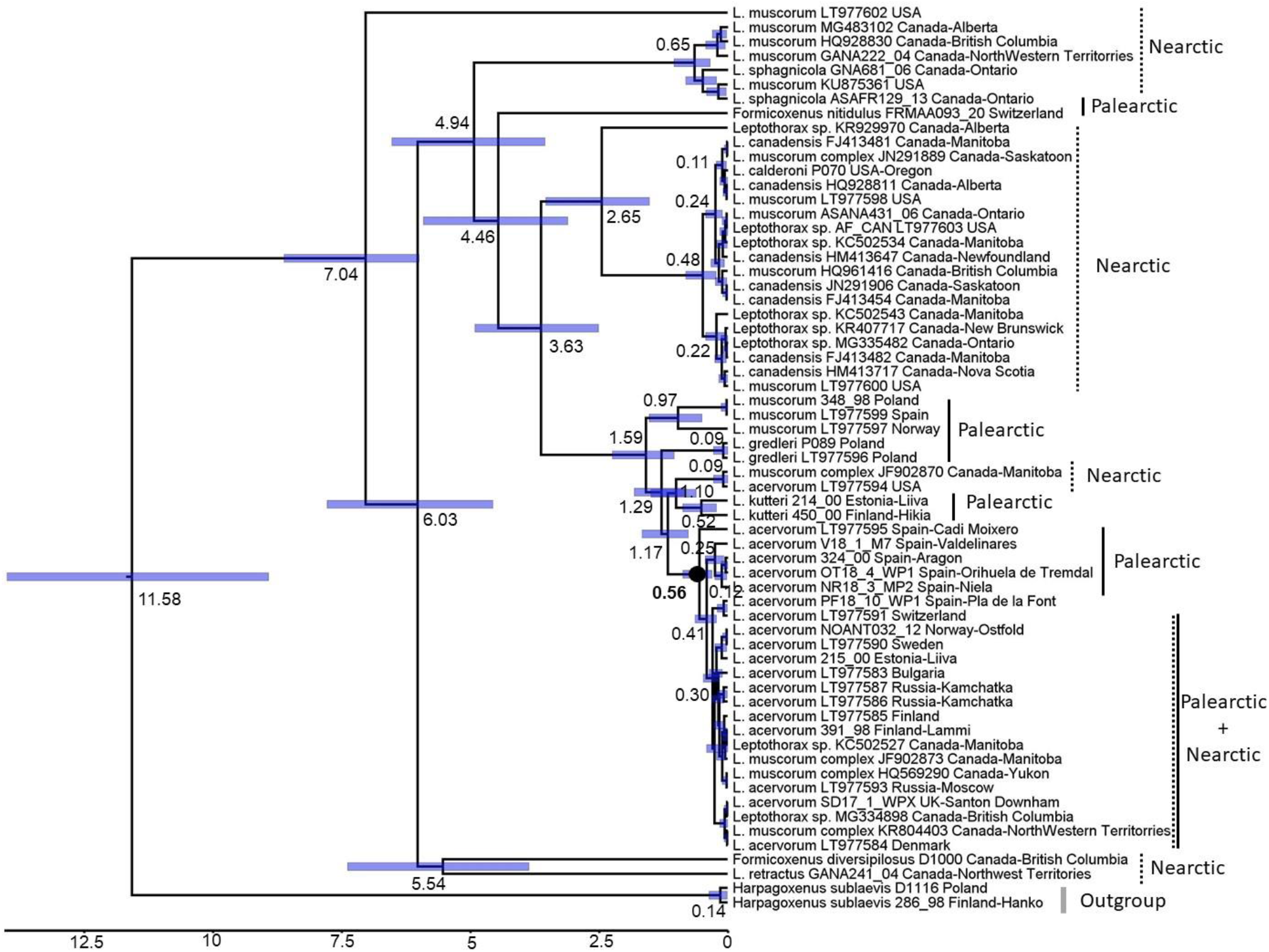
Chronogram of the divergence times estimated in the *Formicoxenus* - *Leptothorax* lineages obtained with Beast. The black circle indicates the monophyletic lineage of *L. acervorum*. Values next to the branches indicate stem ages with blue columns displaying 95% confidence intervals.

**Fig. 4.**
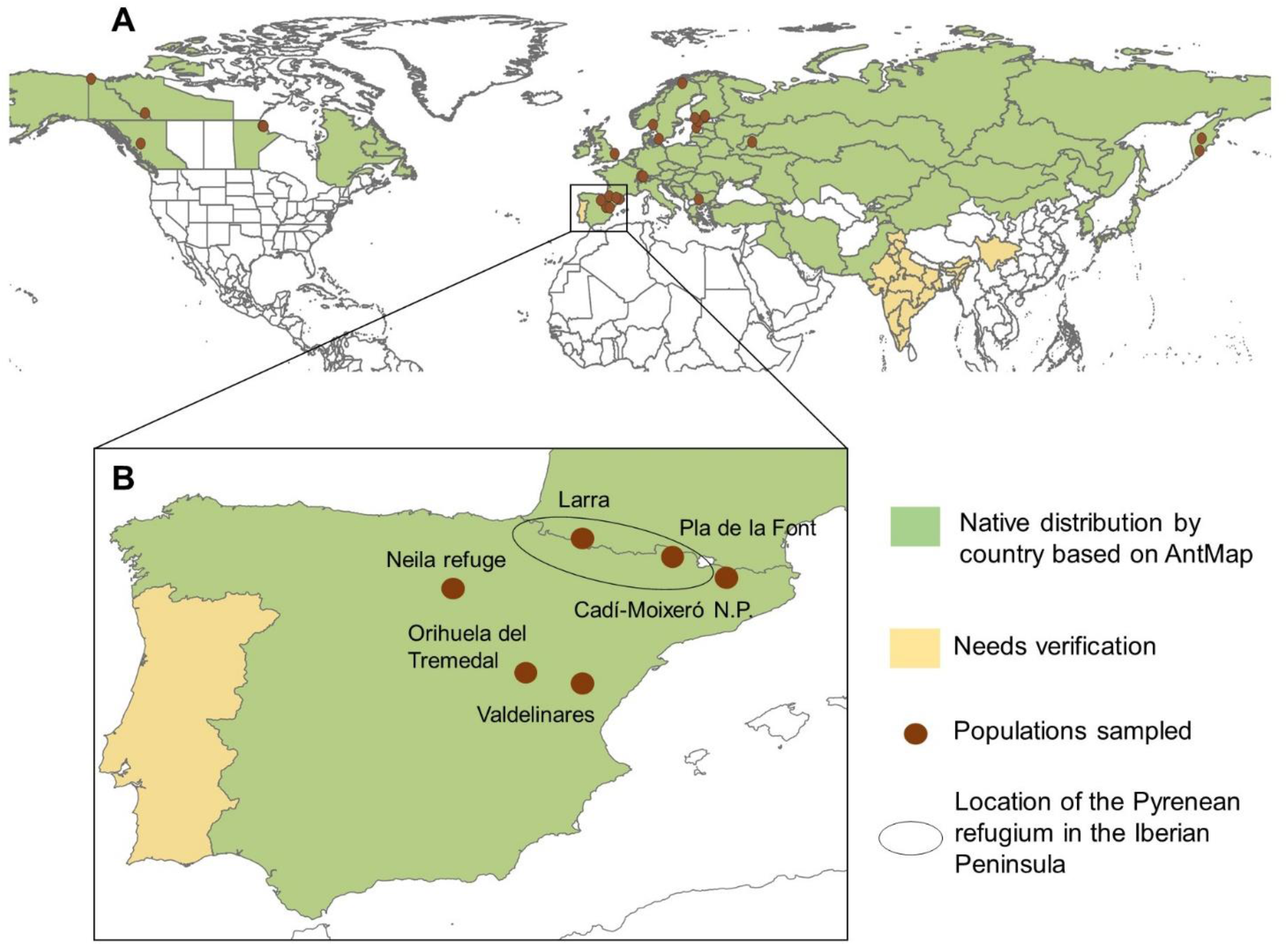
A) Distribution map of *L. acervorum* across the Holarctic region based on the Global Ant Biodiversity Informatics (GABI) database (Guénard et al., 2017). Locations of the different populations included in the phylogeographic analysis are indicated with red dots. B) Close-up of the populations located in the Iberian Peninsula, indicated with blue circles those populations that contributed to the most recent expansion in the Holarctic. The black oval indicates the location of the Pyrenean refugium (Tinaut and Ruano, 2021). N.P. = National Park.

### 3.3 *Phylogeography of* L. acervorum *in the Holarctic region and genetic diversity in the populations of the Iberian Peninsula*

Our analyses based on the COI-5P region (dataset 3, excluding gaps and missing data), recovered 21 variable sites with 19 haplotypes (Hd = 0.826) among the 113 specimens examined (Table S3). Only two populations from the Iberian Peninsula (Larra and Pla de la Font) shared haplotypes with the rest of the populations in the West Palearctic, East Palearctic and the Nearctic. We also found a unique haplotype (H4) shared between the population in England and Switzerland, and the presence of unique haplotypes in Bulgaria and Kamchatka. The most widely distributed haplotype (H11) was shared across the entire geographic distributional range (Table 1), and it likely represents the most recent expansion across the distribution of *L. acervorum*. The haplotype network indicates that most populations in the Iberian Peninsula have been isolated from the remaining range of distribution in the West Palearctic, with a recent expansion of the haplotype H11 into this region (Fig. 5). Our examination of the genetic diversity in the populations in the Iberian Peninsula based on a larger segment of the mitochondrial COI-*trnL2*-COII (dataset 4) suggests higher genetic diversity in Niela refuge and Larra than in the other Iberian populations (Table 2).

**Table 1.**
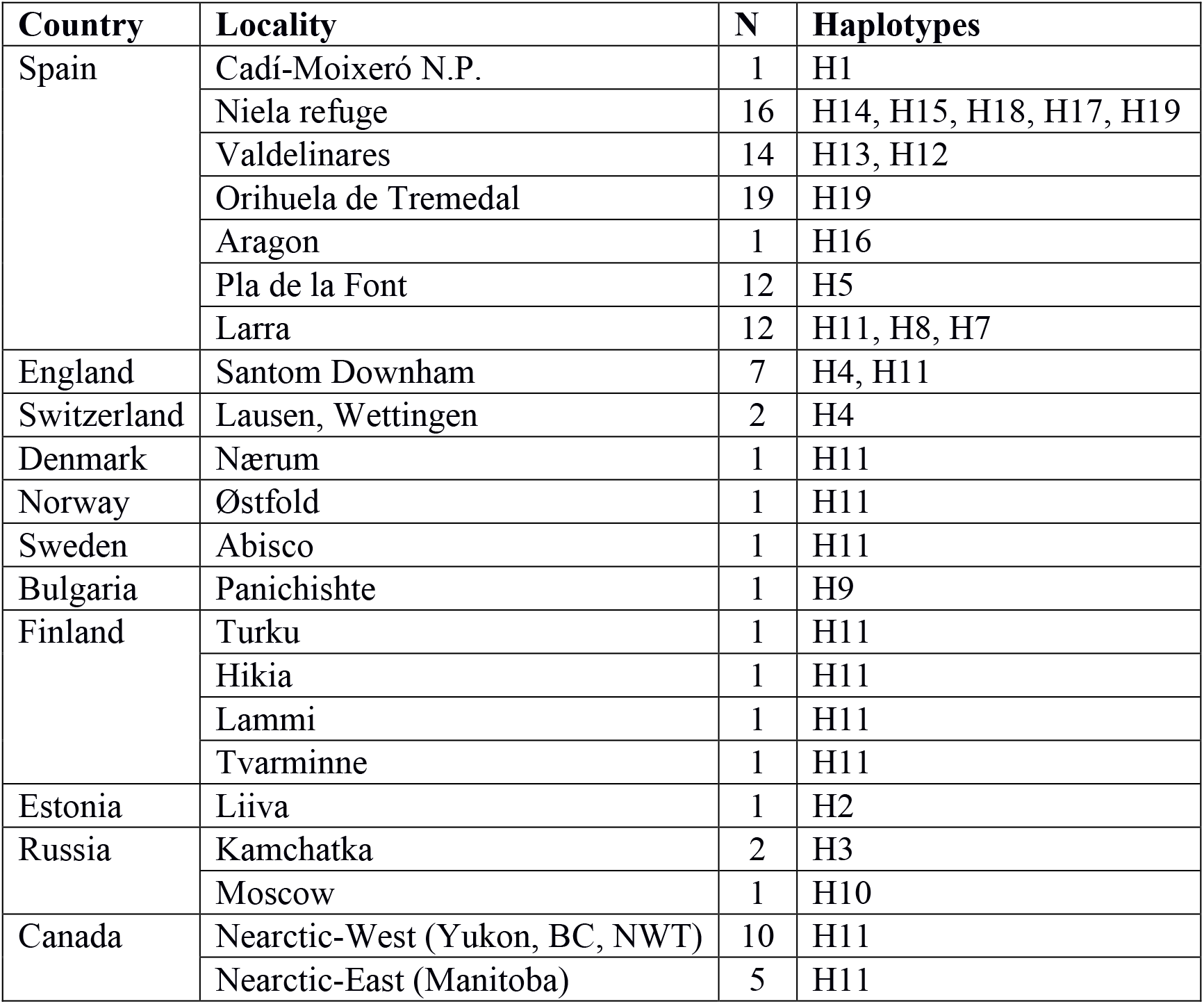
Details of the sampling localities included in the phylogeographic analysis of *L. acervorum* across its Holarctic distribution and the haplotypes observed in each population. N = number of individuals in each locality, BC = British Columbia, NWT = Northwest Territories.

**Table 2.** Estimates of genetic diversity and neutrality tests obtained in the seven populations of *L. acervorum* analyzed in detail.

| Population | N | Ncol | Polymorphic sites | PI | No. haplotyes | *Hd* | $\pi$ | Tajima's D | Fu's *F* |
| --- | --- | --- | --- | --- | --- | --- | --- | --- | --- |
| **Spain** |  |  |  |  |  |  |  |  |  |
| Larra | 12 | 7 | 5 | 3 | 6 | 0.803 | 0.00072 | -0.0093 | -1.847 |
| Niela refuge | 16 | 13 | 16 | 15 | 6 | 0.817 | 0.00228 | 0.3364 | 2.118 |
| Orihuela de Tremedal | 19 | 11 | 8 | 0 | 2 | 0.105 | 0.00037 | -2.1619 | 2.452 |
| Pla de la Font | 12 | 11 | 0 | 0 | - | - | 0 | - | - |
| Valdelinares | 14 | 12 | 1 | 1 | 2 | 0.363 | 0.00016 | 0.3244 | 0.643 |
| **England** |  |  |  |  |  |  |  |  |  |
| Santon Dunham | 7 | 3 | 15 | 14 | 3 | 0.600 | 0.00340 | 1.1546 | 4.116 |

**Fig. 5.**
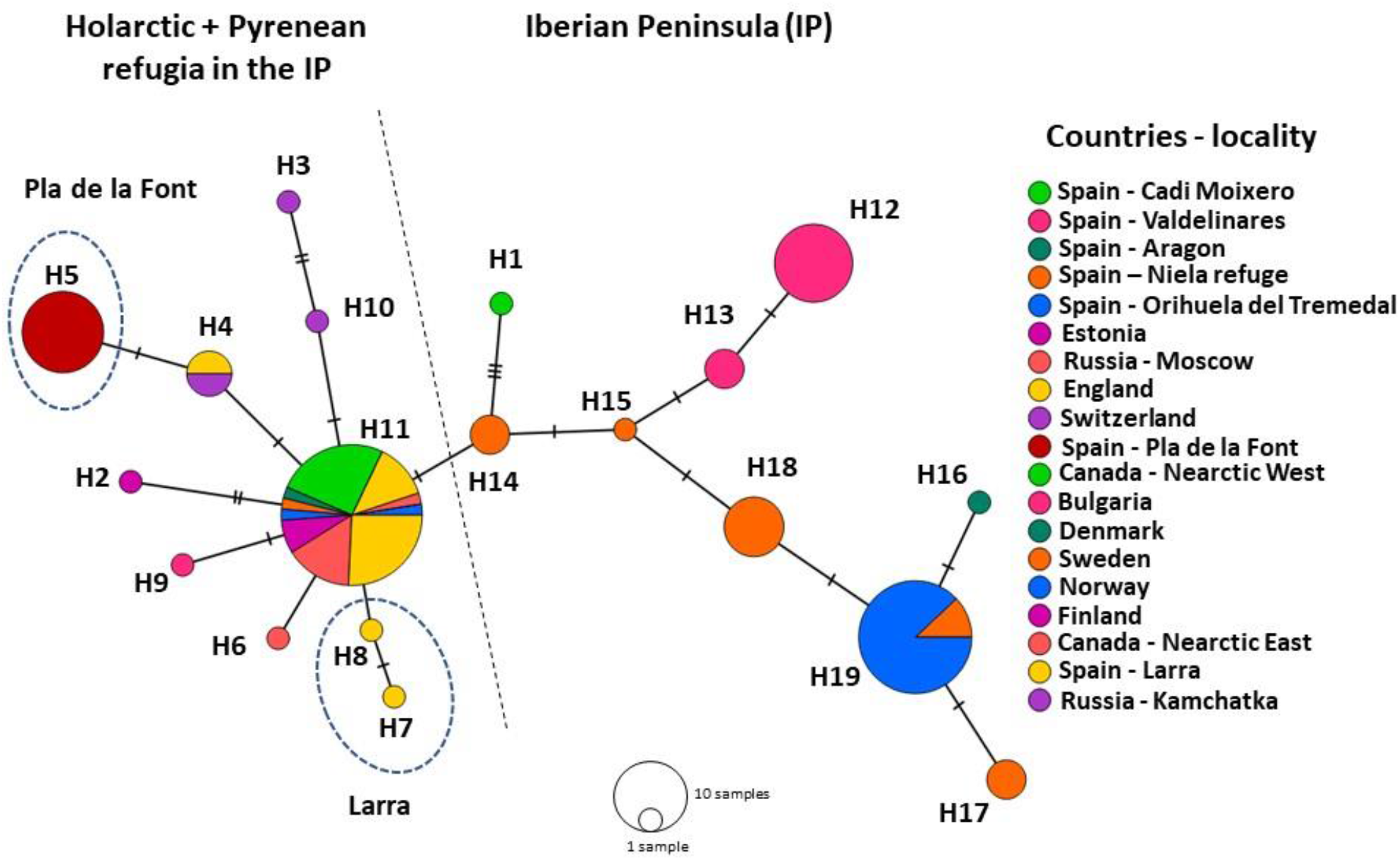
Haplotype network of *L. acervorum* across its Holarctic distribution range. Codes next to the circle indicate the haplotype classification and their distribution. Hatch marks represent mutation differences among the haplogroups. The dash line separates most of the Iberian Peninsula populations from the rest of distribution. Pla de la Font and Larra (in dash circles) are both located in the Pyrenean refugia.

## 4. Discussion

### 4.1 Congruence between phylogenomic inferences based on mitochondrial genomes and UCEs

Recent analyses using large sequence datasets from UCEs have been employed to resolve relationships among ant subfamilies (Blaimer et al., 2015; Branstetter et al., 2017; Li et al., 2018; Longino and Branstetter, 2021). Phylogenomic inferences using mitochondrial genomes have also been used in several ant subfamilies with mostly congruent topologies between nuclear (UCEs) and mitochondrial genomes (Allio et al., 2020). Although we did not have a comprehensive mitochondrial representation within each of the six tribes currently recognized within Myrmicinae (Borowiec et al., 2020), the topology we recovered from this analysis (Fig. 1) is congruent with the known relationships among these tribes inferred from nuclear genes (Ward et al., 2015; https://antwiki.org/wiki/Phylogeny_of_Myrmicinae). Crematogastrini harbors 40% of all Myrmicinae species and 45% of the genera belong to this tribe (Blaimer et al., 2018). Recent phylogenomic analyses based on UCEs (Blaimer et al., 2018; Prebus, 2017), as well as mitochondrial genomes (Park et al., 2021) have been used to increase resolution among genera of of Crematogastrini. Analyses using UCEs have recovered eight clades with high support within Crematogastrini, which have been treated as informal genus-groups.

The *Formicoxenus* genus group consists of six genera (*Formicoxenus, Leptothorax, Vombisidris, Gauromyrmex, Harpagoxenus* and *Temnothorax*) and the relationships among the genera are relatively well established (Blaimer et al., 2018; Prebus, 2017), but less attention has been paid to the *Leptothorax* genus group, which consists of *Formicoxenus, Leptothorax* and *Harpagoxenus*. Species within these genera are prone to develop social parasitism, a set of interrelated lifestyles where the parasitic species depend upon a free-living host to complete their life cycle (Beibl et al., 2005; Heinze, 1995). These three genera, together with *Temnothorax*, are considered a hot spot for the evolution of social parasitism, where it has evolved at least five times among closely related taxa (Beibl et al., 2005; Jongepier et al., 2021; Prebus, 2017).

*Formicoxenus* comprises about seven species of Nearctic or Palearctic distribution, with some species (“guest ants”) living in the nest of *Formica* Linnaeus, 1758, *Myrmica* Latreille, 1804 or *Manica* Jurine, 1807 species. Workers of *Formicoxenus* species beg for food from their host species, but rear their own brood in own chambers within the host nest (Franceur et al., 1985; Heinze, 1995). *Leptothorax* contains some species that are workless social parasites that tolerate (inquilines, e.g., *L. kutteri, L. pacis* (Kutter, 1945) or kill the host queen (murder parasites, e.g., *L. goesswaldi* Kutter, 1967, *L. wilsoni* Heinze, 1989) of other closely related *Leptothorax* species (Foitzik et al., 2009; Heinze and Ortius, 1991). In contrast, *Harpagoxenus* taxa are slave-making (dulotic) ants whose queens kill or expel all adult residents after invading *Leptothorax* spp. (Brandt et al., 2007; Fischer-Blass et al., 2006; Heinze and Ortius, 1991; Pusch et al., 2006).

### 4.2 *Evidence on non-monophyletic taxa within* Leptothorax *and monophyletic lineage within* L. acervorum

Mitochondrial genes (cytochrome b and cytochrome oxidase subunit 1) alone or in combination with other nuclear markers have been previously used in phylogenetic inferences in *Leptothorax* (Baur et al., 1996, 1995; Beibl et al., 2005; Heinze and Gratiashvili, 2015; Schär et al., 2018), but with limited representation of its species or without using other genera within the *Formicoxenus* genus-group. Our more comprehensive sampling (42% of *Leptothorax* species, AntWeb ver. 8.42, https://www.antweb.org, accessed 29 October 2020 and multiple accessions of *L. acervorum*) supports a close relationship between *Formicoxenus* and *Leptothorax* (Fig. 2), similar to previous analyses based on limited taxa of this group (Blaimer et al., 2018; Prebus, 2017; Ward et al., 2015). As it has been previously suggested (Heinze and Gratiashvili, 2015; Heinze and Ortius, 1991; Schär et al., 2018), some taxa within the genus *Leptothorax*, particularly the Nearctic ones, represent species groups that deserve taxonomic adjustments. Our analyses suggest the presence of at least four Nearctic lineages (monophyletic groups with moderate to high support) comprising taxa currently assigned to *L. muscorum* (Nearctic), *L. canadensis, L. calderoni, L. sphagnicola* and specimens assigned to the *L. muscorum* complex. This latter complex seems to consists of a species group of three to four different taxa from the Nearctic that display a set of similar morphological characters and chromosome numbers (Brown, 1955; Heinze, 1991, 1989; Loiselle et al., 1990). One of these lineages also includes a specimen of *L. acervorum* (LT977594), which is more closely related to the Palearctic taxa *L. muscorum, L. gredleri* and *L. kutteri* (Fig. 2). This latter lineage deserves further exploration, as it involves determining whether *L. acervorum* in the Nearctic represents a separate lineage from the remaining samples we included.

In contrast, the Palearctic species *L. gredleri, L. muscorum* and the inquiline *L. kutteri* most likely represent monophyletic lineages (Fig. 2). The lineage of *L. acervorum* comprises both specimens from the Palearctic and Nearctic regions and our divergence age estimates suggest that this clade likely represents the most recent diversification event, within the last 0.5 Ma (Fig. 3). Despite the high support values observed for most of the lineages in the Palearctic region, the support values for the mutual relationships of the lineages were low, and therefore more informative regions will be necessary to determine their relationships.

### 4.3 *Evidence of isolated populations in the Iberian Peninsula with limited contribution to the most recent expansion of* L. acervorum

*Leptothorax acervorum* is one of the only three ant species with Holarctic distribution (Schär et al., 2018); however, all current phylogeographic analyses of this species based on alloenzymes, SSRs, mtDNA and nuclear markers have included only Palearctic specimens (Brandt et al., 2007; Foitzik et al., 2009; Gill et al., 2009; Heinze et al., 1994; Stille and Stille, 1993; Trettin et al., 2016), thus providing an incomplete picture of the patterns of recolonization and populations structure. Previous phylogeographic analyses within *L. acervorum* species have been based on the COI 3’P region and indicate substantial genetic diversity within the species (Brandt et al., 2007; Foitzik et al., 2009; Trettin et al., 2016). The most recent analyses based on SSRs and mtDNA (COI 3’P region) have found the existence of multiple refugia in SW-Europe, and evidence of spatial genetic structure across the sampled area (Trettin et al., 2016). Our most extensive data set, including specimens we identified in the previous phylogenetic analyses (Fig. 2) from the Nearctic region and several populations from the Iberian Peninsula (IP), suggests that the IP populations represent the less derived lineages and that they might have experienced fragmentation and isolation from the remaining Holarctic distribution (Fig. 5). *Leptothorax acervorum* is a cold-adapted species that in the IP inhabits mostly mountainous pinewoods and pine-dominated forest (*Pinus sylvestris*) above 1500 m.a.s.l (Felke and Buschinger, 1999; Gill et al., 2009). Our results indicate that all populations we included within the IP, except Larra and Pla de la Font, seem to have been more isolated from the remaining range of distribution of this species (Fig. 4), supporting previous evidence based on SSRs, which have found evidence of bottlenecks and varying levels of connectivity in this area (Trettin et al., 2016). However, there are only a few mutations separating even the most divergent haplotypes among these populations, but these divergent haplotypes in the IP tend to be found in altitudinally restricted populations. In contrast, the most common haplotype (H11) is found in locations were *L. acervorum* is not altitudinally restricted (Fig. 5) and there is greater connectivity of suitable habitat. The lack of spatial genetic structure previously reported within *L. acervorum* using mtDNA (Brandt et al., 2007; Foitzik et al., 2009; Trettin et al., 2016) might be explained by the limited sampling outside the West-Palearctic regions in previous studies, as well as that this lineage represents the group with the most recent expansion (Fig. 3). Additional sampling across the Holarctic distribution with denser sampling among populations, together with the inclusion of additional markers, would be required to further expand the phylogeographic signal we recovered in our analyses.

Several refugia areas have been identified in the IP based on the ant species in this region (Tinaut and Ruano, 2021), and our results suggests that only populations from the Pyrenean refugia might have more recent connection with the rest of the West Palearctic range of distribution (Fig. 4). In contrast, the populations located in the Cantabric and the Northern Plateu (Tinaut and Ruano, 2021) were likely more isolated from the rest of the populations. Cold-adapted species (boreal) with wide distribution in the Palearctic could survive in periglacial areas during the periods of maximum glacial expansion (e.g., during the LGM, 23-18 ka BP), expanding their range into southern areas. During periods of postglacial warming, southern populations of these species became isolated in mountainous regions (Schmitt, 2009; Schmitt and Varga, 2012), surviving in southern refugia (Stewart et al., 2010). There is extensive evidence of the glacial-interglacial cycles during the Quaternary having influenced the individual genetic diversity and population structure of plants and animals in the West Palearctic (Bennett et al., 1991; Morales-Barbero et al., 2018; Schmitt and Varga, 2012; Stewart et al., 2010), including the presence of several periglacial and southern refugia of cold-tolerant of *Myrmica* (Leppänen et al., 2013, 2011) and *Formica* ant species (Goropashnaya et al., 2007, 2004). Emerging evidence seems to indicate that these glacial-interglacial cycles could also have shared refugia; for example, the congruent phylogeographic signal between *Myrmica* ants and *Betula* species (Leppänen et al., 2011; Maliouchenko et al., 2007), the leaf beetle *Gonioctena intermedia* and its boreal-temperate host trees *Prunus padus* and *Sorbus aucuparia* (Quinzin et al., 2017), and the similar patterns of isolated populations in the Iberian Peninsula observed between *L. acervorum* (Trettin et al., 2016) and *Pinus sylvestris* (Dering et al., 2017; Tyrmi et al., 2020).

## Acknowledgements

The phylogenomic analyses based on whole mitochondrial genomes were performed on resources provided by UNINETT Sigma2 - the National Infrastructure for High Performance Computing and Data Storage in Norway. Funding was provided by NIBIO under the ForGeBiM project. MJ and RLH’s contribution were funded by BBSRC MIBTP PhD studentship to MJ supervised by RLH. We thank Mari Mette Tollefsrud for drawing the map in Fig. 4.

## Author Contribution

DIO, TK, JM and RB conceived the idea; DIO, KV, RS and JM performed the laboratory and analyses; MJ and RLH assembled the *L. acervorum* mtDNA genome, identified the mtDNA genomic scaffold and genotyped re-sequenced samples; DIO wrote the manuscript with contributions from all authors.

## Supporting Information

### Figures

**Fig. S1**. Consensus tree obtained with Bayesian inference as implemented in Mrbayes on the dataset of COI-5P region. The black circle indicates the monophyletic lineage of *L. acervorum* (plus specimens of *L. muscorum* complex and *Leptothorax* sp.) that were later used in the phylogeographic analysis. Values next to the branches represent Bayesian support.

### Tables

**Table S1**. List of specimens used in the phylogenomic analysis of the *Formicoxenus* genus-group using whole mitochondrial genomes. The specimens represent the six genera currently recognized in the group and the outgroup species.

**Table S2**. List of specimens employed in the phylogenetic analyses of *Formicoxenus* - *Leptothorax* using the cytochrome COI-5P region (658 bp).

**Table S3**. Specimens of *L. acervorum* used in the phylogeographic and genetic diversity analyses across its distribution range in the Holarctic region.

**Data S1**. Final alignment of the whole mitochondrial genomes of ant species used in the phylogenomic analysis of the *Formicoxenus* genus-group.

### Supplementary Methods

Methods for the whole mitochondrial genome assembly.

